# Life-history diversity in lakes is associated with ecosystem size

**DOI:** 10.1101/2022.06.01.494213

**Authors:** Andrew L. Rypel

## Abstract

In this study, I inspect how ecosystem size drives fish life-history strategies in north temperate lakes. Species were classified as equilibrium, periodic, or opportunistic strategists according to Winemiller and Rose (1992), and species-area curves assembled and compared among strategy types. The smallest lakes were often dominated by only one species, usually an opportunistic strategist. Overall, species richness rose with ecosystem size, but larger ecosystems tended to be dominated by more periodic and equilibrium strategists. Richness of periodic species increased with ecosystem size at a faster rate compared to opportunistic species. Similarly, life-history niche space increased with ecosystem size in accord with species-area relationships, but showed saturation behavior (i.e., life-history niche space became increasingly ‘packed’ in large lakes). As predicted by theory, relative abundances of opportunistic strategists were extremely variable over space and time, whereas abundances of equilibrium and periodic strategists were more stable. Integration of species-area relationships with life-history theory provides insights into community assembly at multiple scales, and has broad conservation applications.

## Introduction

Ecosystem size is a core driver of species diversity and ecosystem function (MacArthur & Wilson 1967, Post et al. 2000, Fukami 2004). An emerging area of interest is understanding relationships between trait or phylogenetic diversity across islands of varying size or age (Santos et al. 2016, Taylor et al. 2021). Trait-based approaches have a strong foundation in plant community ecology (Kraft et al. 2015, Kimball et al. 2016), but been applied in more limited contexts to other communities (but see Poff et al. 2006, Vaughn et al. 2007, Luiz et al. 2019). One critique of such approaches is that ‘traits’ can be broadly defined, e.g., using available data and idiosyncratic taxa or studies. Further, many traits are collinear, making it difficult to disentangle colonization, extinction and metapopulation dynamics (Tonkin et al. 2018).

Life-history theory provides strong and long-standing frameworks for contextualizing species-area relationships. Winemiller and Rose (1992) identified three strategies in fishes that derive from evolutionary trade-offs among demographic parameters. ‘Equilibrium’ species (essentially the classic ‘K’ strategy described by Pianka (1970)), are characterized by late maturation, low-to-moderate clutch size, and high survival at early life stages (Winemiller 2005, Mims & Olden 2012). ‘Periodic’ strategists have large body sizes, delayed maturation, high fecundity and low juvenile survivorship (Winemiller 1992, Rypel et al. 2021, Tonkin et al. 2021). ‘Opportunistic’ species are small-bodied, fast maturing species with high intrinsic rates of population increase (Xiang et al. 2021). This framework has been key to advancing the field of functional ecology, with application to a range of complicated questions (Haag 2012, Darling et al. 2013, Röpke et al. 2017, Arantes et al. 2019, Malik et al. 2020, Mouillot et al. 2021).

Lakes provide intriguing case studies for how ecosystem size might shape diversity of life-history strategies. For some time, lakes have been recognized as the freshwater equivalent of islands, making them a bridge in application of island biogeography theory to other system types (Eadie et al. 1986, Magnuson et al. 1998, Arnott et al. 2006). Yet lakes may be more isolated than ocean islands, and thus more difficult to colonize (Barbour & Brown 1974). Because fishes express a high degree of diversity in critical demographic parameters (Winemiller & Rose 1992), they also function as excellent case studies for how environmental variations trigger trade-offs in critical demographic rates that might in turn drive community assembly.

The goals of this study are to 1) classify life-history strategies of freshwater fishes in north temperate lakes; 2) quantify and compare species-area relationships across the three broad life-history strategist groups; 3) explore the relationship between species life-history niche space and ecosystem size; and 4) test whether spatial and temporal heterogeneity in species abundances is linked to life-history strategy.

## Methods

The 40 study lakes are based in the north temperate region of USA, mostly within Vilas County, Wisconsin. This region represents one of the densest lake districts in the world and hosts the North Temperate Lakes Long-Term Ecological Research Program (NTL-LTER). A substantial fraction of information presented in this study derives from publically available NTL-LTER data collected over the last 42 y. I also attempt to marry NTL-LTER data with fish assemblage data from two earlier studies in the same region (Tonn & Magnuson 1982, Tonn et al. 1983).

For NTL-LTER, whole fish communities have been sampled annually and consistently since 1981 in seven lakes (Allequash, Big Muskellunge, Crystal Bog, Crystal Lake, Sparkling Lake, Trout Lake, Trout Bog) and since 1995 in four additional lakes (Fish, Mendota, Monona, Wingra). In general, sampling is at the same sites/habitats within lakes over time. Populations are sampled annually with a variety of standard gear types (i.e., fyke nets, gill nets, minnow traps, seines, electrofishing) and all fishes counted. Species richness in each LTER lake was indexed as the count of fish species sampled over the full 42 y period of record in each lake. This approach may include species that locally immigrated or were extirpated over this time frame; however given the imperfect nature of the fisheries gears (Magnuson et al. 1994) it is simply not plausible to accurately track colonization and extirpations of rare fishes. NTL LTER lakes generally represent larger lakes in the landscape with median area = 136.2 ha (range: 1.0-3961.2 ha).

Because NTL-LTER lakes are biased towards large lakes, data were further supplemented with community fish data for small-to-medium sized lakes in the same region using Tonn and Magnuson (1982) and Tonn et al. (1983). Sampling methods are thoroughly summarized in these papers, briefly however, fishes were sampled in 29 lakes with median area = 9.3 ha (range: 0.8-89.8 ha). Fish communities in each lake were intensively sampled during winter and summer using minnow traps and small fyke nets and summer sampling also included a trammel net. Summer and winter samples were pooled to index overall richness in each lake. Lake volumes were estimated as the product of lake area x mean depth for most lakes. For any lakes missing mean depths (n=2 lakes), volume was estimated as a hyperbolic sinusoid (0.43 x area x maximum depth), following Post et al. (2000).

Life-histories of all sampled species were classified according to the Equilibrium-Periodic-Opportunistic (E-P-O) triangular continuum of life history strategies of Winemiller and Rose (1992). The same database of teleost life-history traits used in Winemiller and Rose (1992) was also used in this study (K.O. Winemiller, unpublished data). However, data were enhanced by also including species from the study lakes that did not overlap with the original Winemiller and Rose (1992) work (Supplementary Dataset 1). The final life-history database contained life-history information on 259 fishes, including all 71 sampled species. Parallel to Winemiller and Rose (1992) a Principal Components Analysis (PCA) was conducted on log_10_(age-at-maturity), log_10_(fecundity), and log_10_(juvenile survivorship) data. Juvenile survivorship was defined as the sum of a parental care score described in Winemiller (1989) and average ovum diameter (Winemiller & Rose 1992). K-means cluster analysis (constrained to produce three clusters) was performed on the two principal component scores, which combined to explain 93% of life-history variance in the PCA, to generate an E-P-O triangular classification system of life history strategies (Supplementary Dataset 1). Whereas many species occupy intermediate positions between classes, a soft classification was also built within each class based directly on Olden and Kennard (2010) and Mims and Olden (2013). Briefly, Euclidean distances between each species and its class endmember were calculated such any species could be ranked or weighted according to strategy affinity (Supplemental Dataset 2).

In each lake, species richness and intra-strategy richness (i.e., number of equilibrium, periodic or opportunistic species) were tabulated (Supplementary Dataset 3). Log_10_(richness) and log_10_(strategy richness) were then modeled as functions of log_10_(ecosystem size) using Analysis of Covariance (ANCOVA). In the models, log_10_(richness) was the dependent variable, log_10_(ecosystem size) was the independent variable, and life-history strategy was a blocking variable. Differences in species-area relationships (i.e., slopes or ‘z-scores’) were assessed via log_10_(ecosystem size) x strategy interaction terms.

Because species richness is a coarse measure of species packing within each life history strategy or ecosystem, a life-history niche measure was also devised. Here, I deploy a measure of convex hull area of PCA bi-plot space encompassed by all species, both within a lake overall, and within a given life-history strategy. Use of convex hulls in ecology are discussed in detail in Cornwell (2006). In short, a convex hull functionally outlines the smallest convex set of points containing the same or larger set of points. Hull areas therefore measure total amount of life-history niche space overall, and by strategy, within a given lake. Similar measures have been deployed in other life-history studies (Winemiller et al. 2015, Pianka et al. 2017, Pelegrin et al. 2021); however this study most closely parallels Layman et al. (2007) for food web niche space.

A combination of ANCOVA, breakpoint analysis and logistic regression was used to investigate how life-history niche space scales with ecosystem size. Parallel to richness-ecosystem size analyses, ANCOVA was also used to test relationships between niche space and ecosystem size. In models, log_10_(life-history niche space) was the dependent variable, log_10_(ecosystem size) was the independent variable, and strategy was a blocking variable.

Differences in niche space-area relationships across strategies were again assessed via the log_10_(ecosystem size) x strategy interaction term. However, it was also apparent from plots and residuals that relationships might be non-linear, where rate of niche volume increase declined in larger ecosystems. To better explore threshold values for these shifts, breakpoint regressions were developed using program SegReg. Breakpoint analysis asks the question whether a given data series is better defined by one line or two. If two, the analysis will estimate functions for both lines in addition to a threshold or ‘breakpoint’ value where two lines diverge. Finally, because communities of organisms can also be considered fractionally, proportion of species belonging to each life-history strategy in each lake was also calculated. To estimate proportional niche space, convex hull area of each strategy was calculated as above and divided by total convex hull area for all species in a lake. Proportional niche space values were then plotted against ecosystem size, and a logit-based logistic regression used to test the relationship.

A key prediction emerging from Winemiller and Rose (1992) is that spatiotemporal variability in population numbers should also have a basis in life-history strategies. Opportunistic species are predicted to have high spatiotemporal heterogeneity in abundance as an adaptation to unstable environments. In contrast, equilibrium species are predicted to have low spatiotemporal abundance heterogeneity because of evolution towards nesting and nest guarding, which necessitate environmental stability. Spatial-temporal abundance heterogeneity data are based on Magnuson et al. (2021) but summarized for this analysis in Rypel (2021). Spatial and temporal abundance heterogeneity was indexed as the coefficient of variation for each species in CPUE:

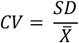

where CV is the coefficient of variation of the sample, SD is the sample standard deviation, and 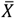 is the sample mean. Differences among strategies in spatial and temporal abundance heterogeneity were tested using weighted Analyses of Variance (weighted ANOVAs). In models, temporal or spatial heterogeneity for each species was a dependent variable, strategy was the independent variable, and strategy affinity from the soft classification was a weighting factor.

## Results

71 fish species were captured over space and time across all lakes (Figure 1, Supplementary Dataset 1). PCA of log_10_(age-at-maturity), log_10_(fecundity), and log_10_(juvenile survivorship) produced dominant axes (PC1 and PC2) explaining 93% of variance in life-history traits among all fish species. Using k-means cluster analysis on PC1 and PC2 scores, species were classified into opportunistic, periodic and equilibrium life-history strategies, including all 71 north temperate lake species. A total of 16 species (22.5%) were classified as equilibrium, 29 species (40.9%) as opportunistic, and 26 species (36.6%) as periodic. For opportunistic strategists, the three fishes with strongest membership were Iowa darter *Etheostoma exile*, mimic shiner *Notropis volucellus*, and northern redbelly dace *Chrosomus eos*. For equilibrium species, strongest members were yellow bullhead *Ameiurus natalis*, black bullhead *Ameiurus melas*, and slimy sculpin *Cottus cognatus*. For periodic strategists, strongest members were freshwater drum *Aplodinotus grunninens*, bigmouth buffalo *Ictiobus cyprinellus*, and burbot *Lota lota*.

**Figure 1.**
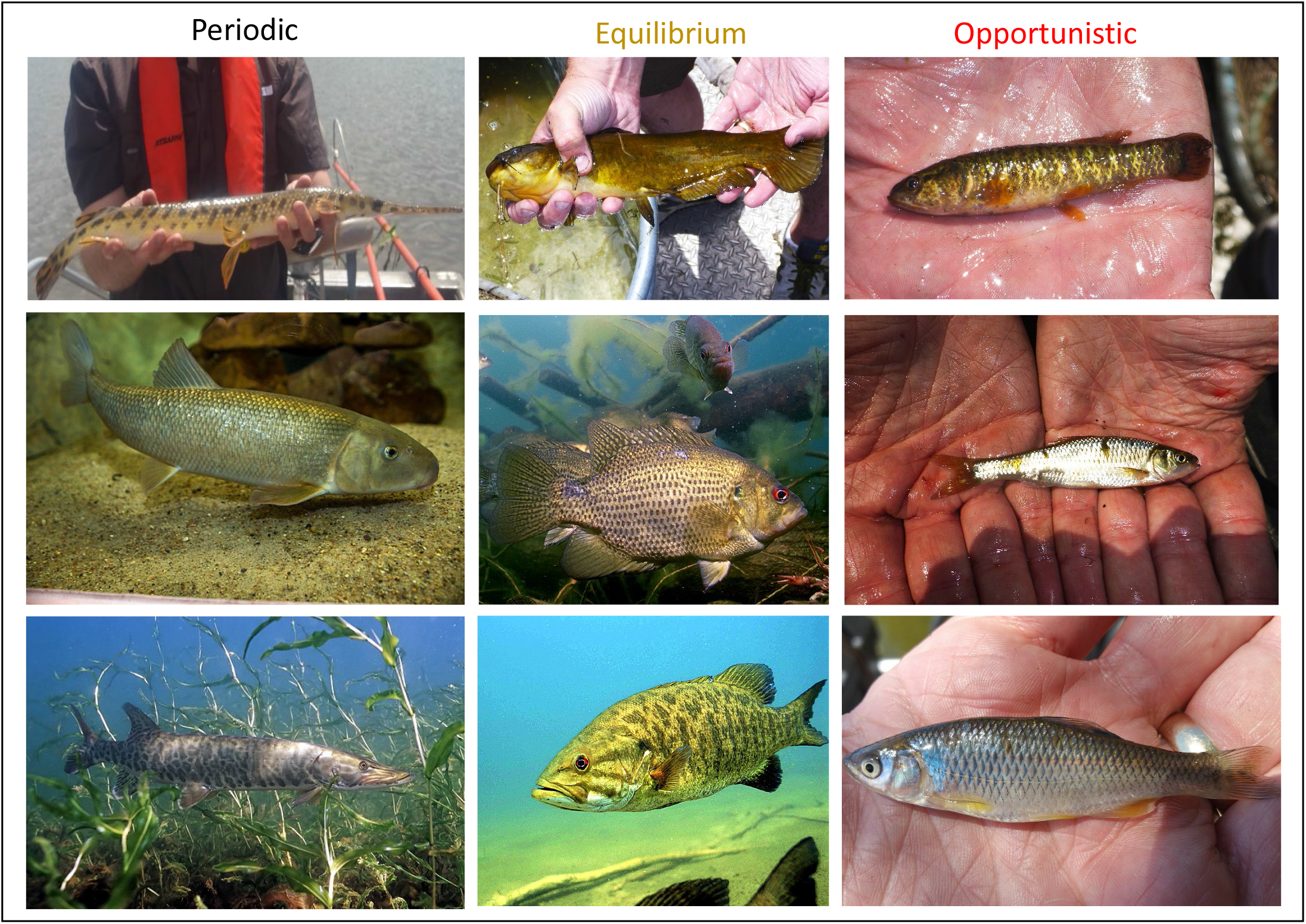
Example study species from each of three life-history strategies in north temperate lakes. Periodic species included (from top to bottom): longnose gar *Lepisosteus osseus*, white sucker *Catostomus commersonii*, and muskellunge *Esox masquinongy*. Equilibrium species included: yellow bullhead *Ameiurus natalis*, rock bass *Ambloplites rupestris*, and smallmouth bass *Micropterus dolomieu*. Opportunistic species included central mudminnow *Umbra limi*, common shiner *Luxilus cornutus*, and spotfin shiner *Cyprinella spiloptera*. Images of white sucker, muskellunge, rock bass, and smallmouth bass all from wikicommons.org. All other images by the author.

Across lakes, overall species richness varied dramatically from 1-45 species. Three small lakes (Trout Bog, Lake 33-13, and Lake 6-7) had just one species. There was a significant relationship between total species richness and lake volume (DF = 38, F = 149.1, P < 0.0001, r^2^ = 0.80, Figure 2). Positive log-linear relationships were also found for intra-strategy richness (DF = 114, F = 44.0, P < 0.0001, r^2^ = 0.66); however slopes and elevations (intercepts) differed significantly by strategy. Periodic richness increased at a faster rate with lake volume than did opportunistic richness (ANCOVA log_10_(ecosystem size) x strategy interaction term, P = 0.04).

**Figure 2.**
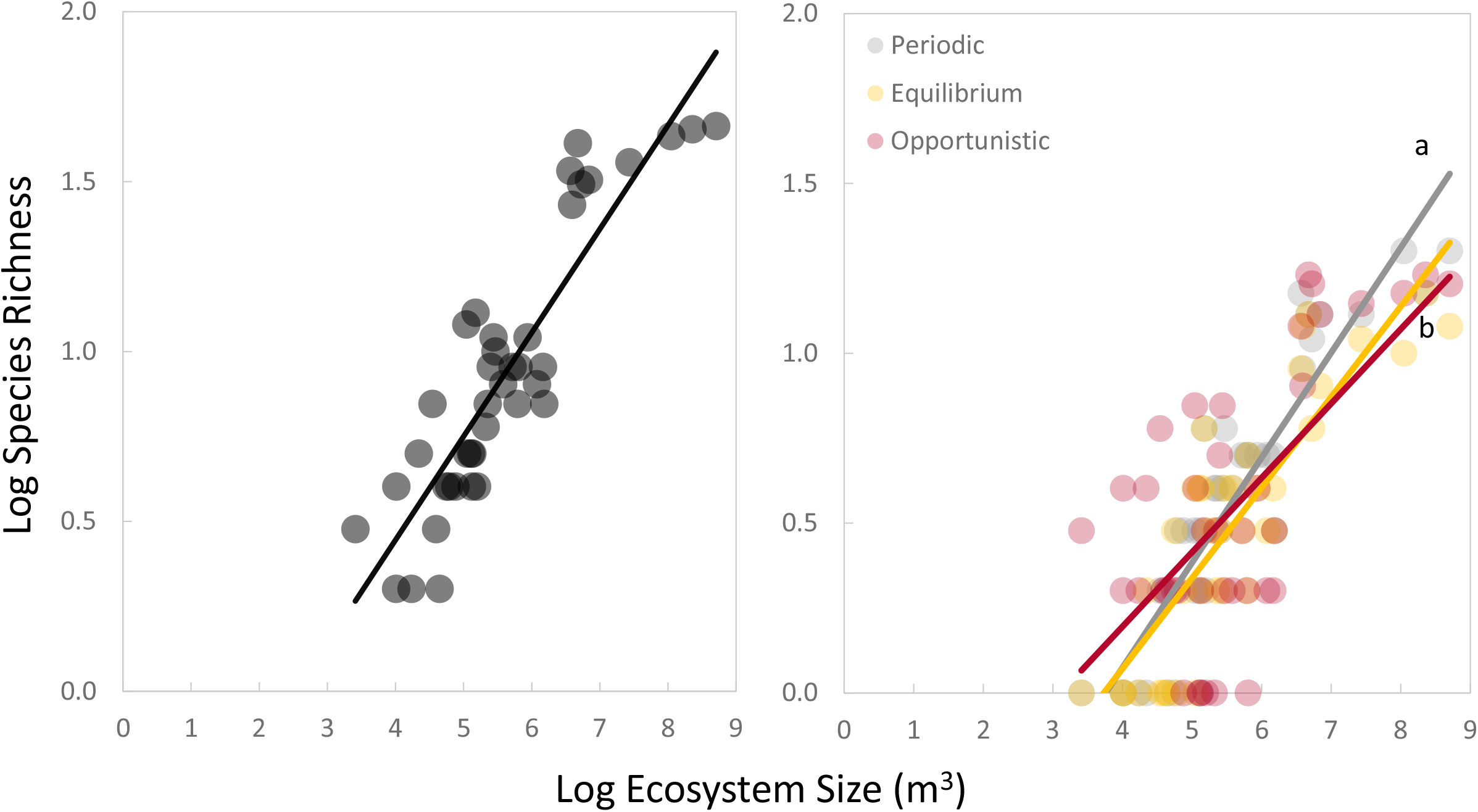
(Left) Relationships between absolute species richness and ecosystem size across 40 WI USA lakes. (Right) Relationships between species richness and ecosystem size for each of three major life-history strategy types. Letters denote slopes that differ significantly.

However, neither periodic nor opportunistic species differed from equilibrium species in slope. Further, no strategies were significant by themselves in the model, indicating mean richness did not differ substantially by strategy. Aside from the interaction term, only volume was a significant predictor of richness (P < 0.0001).

Biplots of PC1 versus PC2 scores were constructed for each lake ecosystem, revealing lake-specific dynamics in strategy diversity (Figure 3). Convex hulls were fit to each lake’s biplot to quantify life-history niche space occupied in each ecosystem in addition to convex hull areas for each of the three strategies. Small lakes with few species had contracted convex hulls (e.g., Bug Lake) relative to moderate-sized (Johnson Lake, Grassy or Crystal Lakes) or larger volume lakes with high richness (e.g., Trout Lake). Furthermore, small lakes were often dominated by opportunistic species. Indeed opportunistic species were the only species that could exclusively dominate communities in the smallest lakes.

**Figure 3.**
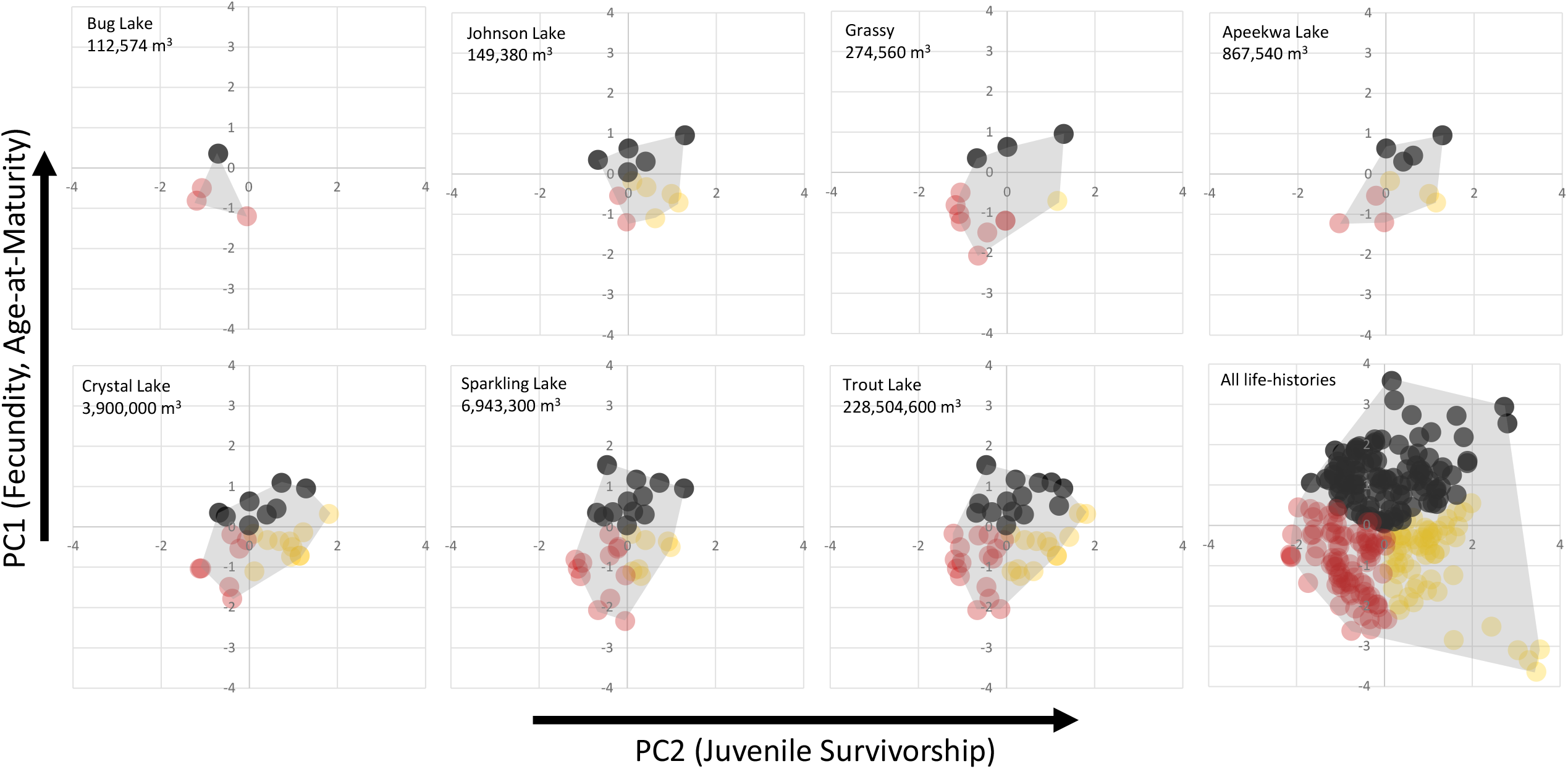
Convex hulls showing examples of how life-history niches enlarge with ecosystem size.

Convex hulls enlarged with lake size for all strategies (Figure 4). When intra-strategy convex hulls were modeled as a function of ecosystem size, a significant pattern was revealed (df = 114 F = 23.0, r^2^ = 0.50, P < 0.0001); however, slopes of relationships again differed by strategy. Periodic species niche space increased at a faster rate with lake volume than did opportunistic species richness (ANCOVA log_10_(ecosystem size) x strategy interaction term, P = 0.03). However, neither periodic nor opportunistic species differed from equilibrium species in slope. In all three cases, data were better described by two lines rather than one (Table 1, Figure 4). For opportunistic species, even though a breakpoint was identified, slopes were best modeled as flat before and after the threshold.

**Table 1.**
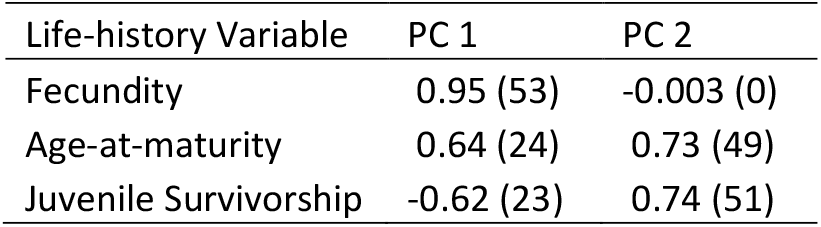
PC loadings and % contribution of variables (in parentheses) extracted from PCA of life-history variables. Eigenvalues were 1.7, 1.1 and 0.2 for PCs 1, 2 and 3, respectively.

| Life-history Variable | PC 1 | PC 2 |
| --- | --- | --- |
| Fecundity | 0.95 (53) | -0.003 (0) |
| Age-at-maturity | 0.64 (24) | 0.73 (49) |
| Juvenile Survivorship | -0.62 (23) | 0.74 (51) |

**Figure 4.**
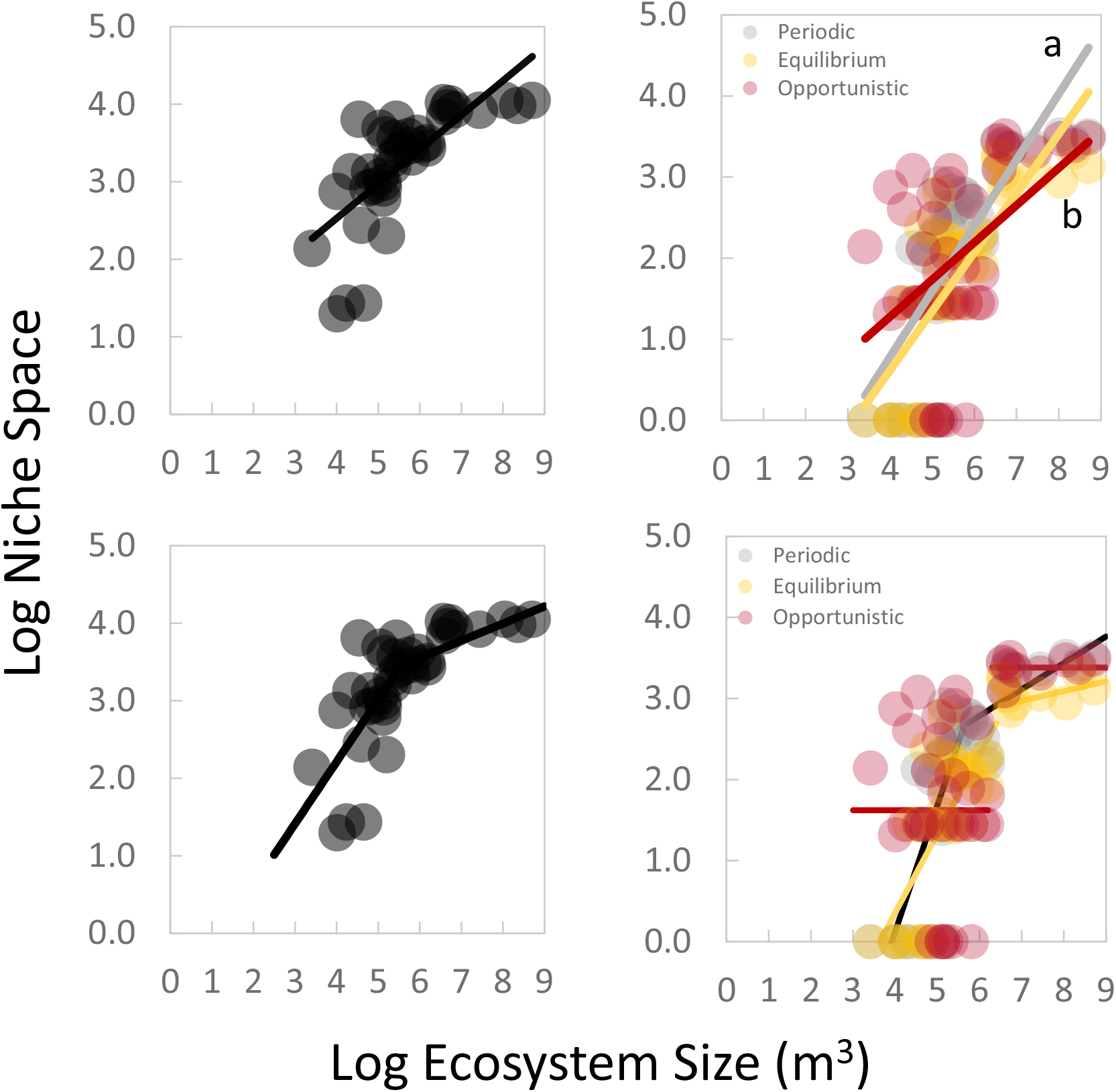
(Upper Left) Log-linear relationship between total convex hull niche space and ecosystem size across 40 lakes. (Upper Right) Log-linear relationships between convex hull niche space and ecosystem size for each of three major life-history strategy types. Letters denote regression slopes that differ significantly. (Lower Left) Breakpoint relationship between total convex hull niche space and ecosystem size. (Lower Right) Breakpoint relationships between convex hull niche space and ecosystem size for each of three major life-history strategy types. Additional results on breakpoint regressions are in Table 2.

Relative proportions of life-history strategies also shifted in response to ecosystem size (Figure 4, right panels). For both periodic and equilibrium species, % niche space increased as a significant logistic function of ecosystem size (logistic regression, model −2 Log(Likelihood) P < 0.0001, R^2^ Nagelkerke = 0.71). In general, % of periodic species increased to an average maximum of ∼30% in the largest lakes. Equilibrium species increased to an average maximum of ∼26% for the largest lakes; however patterns for these species were variable as indicated by their reduced Negelkerke score (logistic regression, model −2 Log(Likelihood) P < 0.0001, R^2^ Nagelkerke = 0.25). Opportunistic species increased to an average maximum of ∼32% of available niche space. And while the logistic model for opportunistic species was also significant, it was the least predictive (logistic regression, model −2 Log(Likelihood) P < 0.0001, R^2^ Nagelkerke = 0.20). Furthermore, opportunistic species were the only life-history that could completely dominate communities in the smallest lakes (100% of niche space).

**Table 2.** Results of breakpoint analyses examining how intra-strategy diversity scales with lake size.

| Life-history | Breakpoint (Y/N) | Breakpoint (log, m<sup>3</sup>) | Eq below breakpoint | Eq above breakpoint |
| --- | --- | --- | --- | --- |
| All | Y | 5.63 | $0.80 \times \log_{10}(\text{ecosystem size}) - 0.98$ | $0.22 \times \log_{10}(\text{ecosystem size}) + 2.23$ |
| Opportunistic | Y | 6.22 | $0.00 \times \log_{10}(\text{ecosystem size}) + 1.62$ | $0.00 \times \log_{10}(\text{ecosystem size}) + 3.38$ |
| Equilibrium | Y | 6.64 | $0.97 \times \log_{10}(\text{ecosystem size}) - 3.54$ | $0.12 \times \log_{10}(\text{ecosystem size}) + 2.13$ |
| Periodic | Y | 5.63 | $1.56 \times \log_{10}(\text{ecosystem size}) - 6.12$ | $0.33 \times \log_{10}(\text{ecosystem size}) + 0.83$ |

Spatial and temporal heterogeneity in fish abundances differed significantly by life-history strategy (Figure 5). For spatial heterogeneity, all three strategies differed significantly in abundance heterogeneity (weighted ANCOVA, df = 3, F = 20.2, P < 0.0001). Opportunistic species had the highest spatial heterogeneity in fish abundance, followed by periodic species, and finally equilibrium species (weighted ANCOVA, LSD post-hoc tests, all P’s < 0.003). For temporal heterogeneity, there were also significant differences in abundance heterogeneity by life-history strategy (weighted ANCOVA, df = 3, F = 20.2, model P < 0.0001). Opportunistic species again had the highest heterogeneity values for temporal abundance heterogeneity, and these values differed significantly from periodic (LSD post-hoc P < 0.0001) and equilibrium (LSD post-hoc P < 0.0001) species. However, periodic and equilibrium species did not differ significantly in temporal abundance heterogeneity (LSD post-hoc P = 0.66).

**Figure 5.**
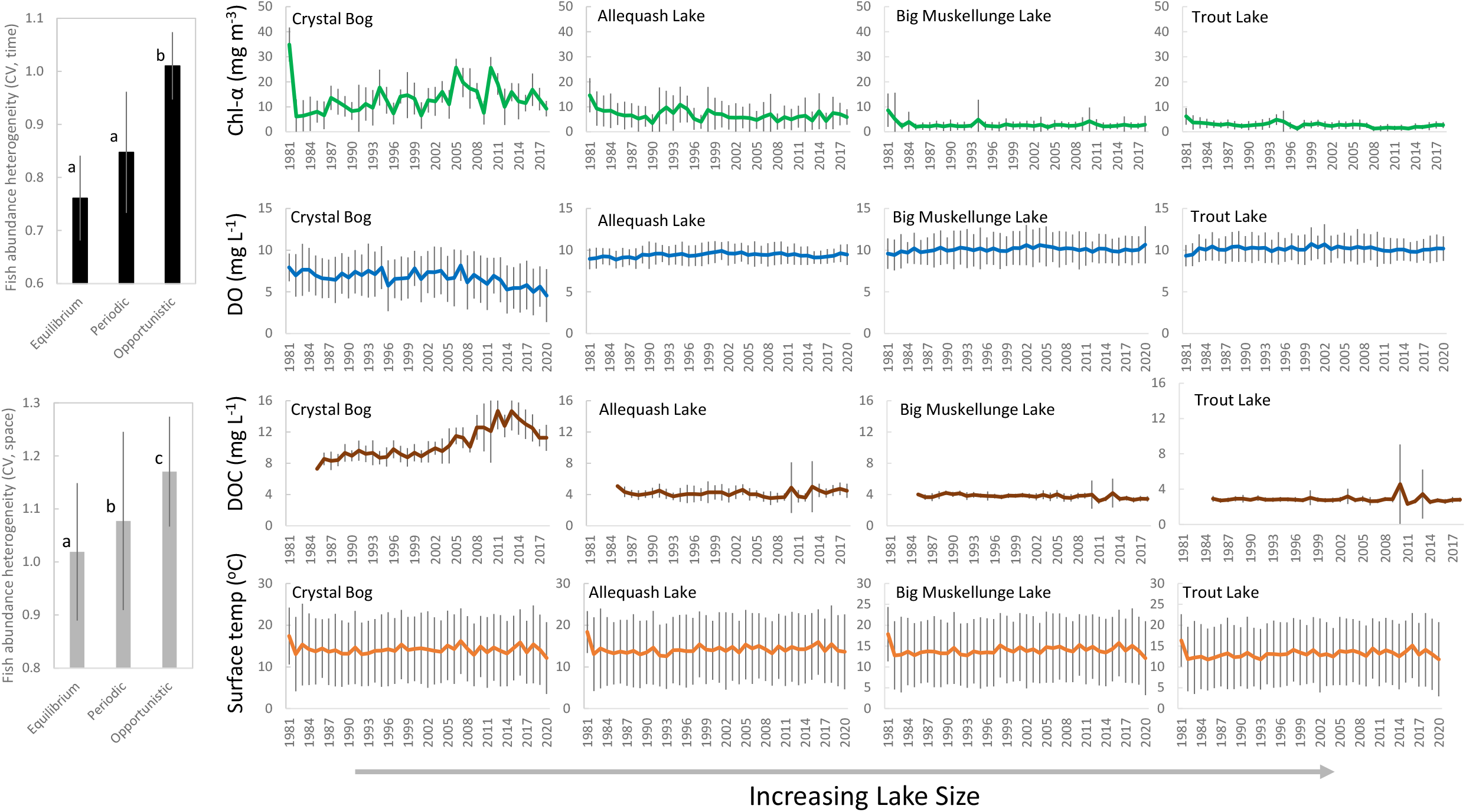
(Left) Comparisons of temporal (top) and spatial (bottom) heterogeneity in fish abundances across three broad life-history strategy groups. Error bars denote the mean ± 1 SE. Letters indicate means that differ significantly using weighted ANOVA. (Right) Plots illustrating long-term variations in critical limnology drivers in four LTER lakes that vary by size. Lines represent annual means and error bars denote the mean ± 1 SE.

## Discussion

This study provides evidence for how life-history and island biogeography are linked ecological concepts. MacArthur and Wilson (1967) famously described an equilibrium point in species-area relationships where colonization approximates extinction. A number of factors influence equilibrium points (e.g., island size), and gear biases in the current study precludes estimation of these points (Magnuson et al. 1994). Nevertheless, it is evident via comparative analyses that life-history diversity in lakes, and species packing dynamics, are associated with ecosystem size.

The z-score (slope of the species area relationship) for all north temperate lakes was high (z = 0.31). In comparison, a large meta-analysis found that z-scores for forests, rivers, islands, streams and lakes averaged 0.34, 0.30, 0.26, 0.21, 0.21, and 0.18, respectively (Drakare et al. 2006). In another study, mean z-score across large lakes globally was just 0.15 (Barbour & Brown 1974). Typically, more isolated islands are considered as having higher z-scores (Preston 1962, Preston 1962). Therefore these north temperate lakes may be more isolated in their community assembly patterns than previously thought.

Ecosystem size appeared to drive patterns of occurrence of all three strategy types: opportunistic species dominated small lakes, periodic species increased in larger lakes, and equilibrium species thrived in intermediate systems. Nonetheless, there were exceptions: opportunistic species accounted for 40.9% of fish diversity overall (versus 22.5% and 36.6% for equilibrium and periodic species, respectively) and were cosmopolitan. That the opportunistic strategy was the most common overall perhaps highlights a heterogeneous element to lakes overall. Results are also congruent with Olden and Kennard (2010) who found opportunistic species were the dominant life-history strategy in streams on both Australian and North American Continents, and that increasing hydrologic variability promoted greater prevalence of opportunistic strategists. Mims and Olden (2012) showed periodic strategists were associated with ecosystems having high seasonality in flows but reduced variability. Moderate-sized lakes appear to encourage higher populations of species that benefit from spatiotemporal stability (i.e. equilibrium strategists). Periodic species seem constrained such that spawning migrations outside the confines of the traditional ‘lake’ are needed to complete complicated life cycles (e.g., into wetlands, stream and large river tributaries). Periodic strategists like sturgeon, walleye, freshwater drum and northern pike all exhibit cross-ecosystem or -habitat migrations (Bruch & Binkowski 2002, Weeks & Hansen 2009, Richard & Rypel 2012) which become more common in larger drainage lakes with increased hydrologic connectivity.

Life-history niche space analyses add additional context to biogeography patterns. More specifically, while overall patterns are similar, they also appear to be non-linear. Thus convex hulls expand with ecosystem size, but this expansion slows after critical points. These findings support an older, but still highly relevant literature on ‘species packing’ (Hutchinson 1959, MacArthur & Levins 1967, Roughgarden & Feldman 1975, Cui et al. 2020), whereby guild diversity stays the same, but number of species per guild or tightness of species packing increases with ecosystem size (Werner 1977, Terborgh et al. 1978, Peay et al. 2007). Brown (1981) noted future attempts to understand species diversity might focus on developing ‘capacity rules’; convex hulls may be a useful tool in this context. When data in this study are compared to ∼6K lakes in Wisconsin known to support fishes (Rypel et al. 2019), breakpoint thresholds were between the 75th-95th quantiles of all known lake volumes in the region. Thus saturation of life-history niche space (i.e., intense species packing) is only occurring in very large lakes.

Nonetheless, that there is a saturation point is itself intriguing. Although immigration rates may be higher in large lakes, there appears to be increased difficulty for new species to establish, perhaps because niches are crowded. Similarly, very small lakes are simply too harsh for the majority of species in the pool. Taken together, analyses demonstrate how life-history strategies modulate colonization and extinction dynamics, which in turn drives commonly-observed community assembly patterns.

Across study lakes, opportunistic species expressed the highest degree of spatial and temporal heterogeneity in abundances (Figure 5). In north temperate lakes, environmental instability is strongly driven by variable physicochemical conditions (Hanson et al. 2008, Fang & Stefan 2009, Bouffard et al. 2012). Indeed, a potential signal is evident that heterogeneity in physicochemistry decreases with lake size (Figure 5) and studying the applicability of this pattern across a larger range of lake sizes might be a question of interest to limnologists. Small lakes are notable for their harshness, including a tendency for fish kills (Magnuson et al. 1985, Fang & Stefan 2000, Till et al. 2019). Regular kills in turn preclude most species from colonizing aside from those with adaptations for colonization and persistence (Tonn & Magnuson 1982), i.e., opportunistic species. Z-scores for periodic, equilibrium, and opportunistic species were 0.31, 0.27, and 0.22, respectively. Thus, opportunistic strategists are the least isolated of the strategies – an apparent evolutionary adaptation to enhance colonization.

These findings have implications for regulation of populations, and conservation management of freshwater ecosystems. For example, management strategies aimed at matching life-history traits to essential habitats could be more effective (Southwood 1977, Winemiller 2005, Sass et al. 2017, Arantes et al. 2019). Using equilibrium species as one example – stable spatial and temporal conditions would likely benefit these species. Management actions that promote an equilibrium context might include hydrological management, i.e., reduced flow variance (Zeug & Winemiller 2008, Rypel 2011, McManamay & Frimpong 2015), shoreline restoration to promote stable and long-lasting spawning and nesting/rearing habitats (Sass et al. 2019), and fishing limits that discourage disruption of reproduction on nests (Cooke et al. 2000). In contrast, periodic species may benefit from a different suite of management decisions. Connectivity of seemingly disparate habitats and ecosystems is likely crucial for the health and balance of these populations (Paukert & Fisher 2001, Rypel et al. 2006, Rypel & Bayne 2009). Therefore, enhancing hydrologic and landscape connectance, in addition to protection and restoration of diverse habitat types would likely aid periodic species conservation (Childress et al. 2014, Pracheil et al. 2019, Rypel 2021).

These findings may be widely applicable, and the E-P-O model has already been applied to freshwater mussels (Haag 2012, Randklev et al. 2016), marine corals (Darling et al. 2013), and even bacteria (Malik et al. 2020). Future work might explore a generality of life history diversity-ecosystem size relationships to more taxonomic groups and ecosystem types. Because evolution selects traits to optimize fitness, it is probable that results do apply in other realms. Further, an approach of building and quantifying convex hulls to index life-history niche space may be attractive, perhaps especially ecologists interested in species packing. Species lists are readily available for a wide array of ecosystem types, and increasingly with modern ecological methods (e.g., environmental DNA).

## Conclusions

Island biogeography and life-history theory are interwoven concepts, but this point, have not been treated as such. Ecosystem size clearly drives patterns in richness, colonization and extinction dynamics of life-histories. There is increasing and wide interest in using species ‘traits’ to understand assembly of diverse communities (Kraft et al. 2008, Laliberté & Legendre 2010, Taylor et al. 2021). However, ‘traits’ may be too broadly defined to be useful in some synthetic constructs. A universal system of quantifying traits might invoke a schema like that of Winemiller and Rose (1992) to generate a common ecological language. For example, life-history of any taxon could theoretically be mapped to the E-P-O triangular continuum. Ultimately, small freshwater ecosystems are likely to be favored by small-bodied species with fast generation times, high juvenile survivorship, and low fecundity (i.e., opportunistic species). In contrast, equilibrium and periodic species are likely to be more common in larger ecosystems with less disturbance and greater connectivity. These fundamental constraints on life-history diversity help clarify the types of organisms that various ecosystems support (Hutchinson 1959), along with appropriate methods for managing or conserving them.

## Acknowledgements

I thank a mass of dedicated scientists who diligently collected scientific information on fishes and lakes at NTL-LTER over the last 50 years. This study was supported by the California Agricultural Experimental Station of the University of California Davis (grant number CA-D-WFB-2467-H). I thank Dr. Kirk O. Winemiller and Dr. John J. Magnuson for reviewing earlier versions of this work. Dr. Winemiller also provided access to the core life-history dataset.

